# TCF7L2 Regulation of GATA6-dependent and -Independent Vascular Smooth Muscle Cell Plasticity and Intimal Hyperplasia

**DOI:** 10.1101/397851

**Authors:** Roshni Srivastava, Harshavardhan Rolyan, Yi Xie, Na Li, Neha Bhat, Lingjuan Hong, Fatemehsadat Esteghamat, Adebowale Adeniran, Arnar Geirsson, Jiasheng Zhang, Guanghao Ge, Marcelo Nobrega, Kathleen A. Martin, Arya Mani

## Abstract

Genetic variations in Wnt-coreceptor LRP6 and Wnt-regulated transcription factor TCF7L2 have been among the strongest genetic signals for type2 diabetes (T2DM) and coronary artery disease (CAD). Mice with a CAD-linked LRP6 mutation exhibit obstructive coronary artery disease characterized by reduced TCF7L2 expression and dedifferentiation of vascular smooth muscle cell (VSMC). While TCF7L2 maintains stemness and promotes proliferation in embryonic tissues and adult stem cells, its role and mechanisms of action in VSMC differentiation is not understood. Using multiple mouse models, we demonstrate here that TCF7L2 promotes differentiation and inhibits proliferation of VSMCs. TCF7L2 accomplishes these effects by stabilization of GATA6 and upregulation of SM-MHC and cell cycle inhibitors. Accordingly, *TCF7L2* haploinsufficient mice exhibited increased susceptibility to, while mice overexpressing TCF7L2 were protected against injury-induced intimal hyperplasia compared to wildtype littermates. Consequently, the overexpression of TCF7L2 in LRP6 mutant mice rescued the injury induced intimal hyperplasia. These novel findings imply cell type-specific functional role of TCF7L2 and provide critical insight into poorly understood mechanisms underlying pathogenesis of intimal hyperplasia.

## Introduction

Advances in medical therapy such as widespread use of lipid lowering agents led to a significant decline in prevalence of coronary artery disease (CAD) mortality over previous four decades. The recent rise in cardiovascular death has generated renewed interest in identifying novel disease risk factors (Martin-Timon, Sevillano-Collantes et al., 2014). A parallel rise is observed in prevalence of T2DM and obesity, which markedly increases the risk of myocardial infarction and restenosis. The pathogenesis of T2DM-linked CAD is most likely distinct from those extensively studied in animal models of severe hyperlipidemia and inflammation, necessitating new disease models. Recent studies, including GWAS of CAD and myocardial infarction (MI) have implicated altered differentiation of VSMC in pathogenesis of CAD (Motterle, Pu et al., 2012, Nurnberg, Cheng et al., 2015). VSMC are the most abundant cell type and have the highest degree of plasticity among all vascular cells (Majesky, Dong et al., 2012, Majesky, Dong et al., 2011, Rensen, Doevendans et al., 2007). The phenotypic switching of VSMC allows their migration into the intima where they continue proliferating and depositing extracellular matrix (Owens, Kumar et al., 2004) or transdifferentiate into other cell types such as CD-68 positive and foam cells (Shankman, Gomez et al., 2015). Whether VSMC proliferation has a beneficial or detrimental role in atherosclerosis process is controversial. VSMCs may have a protective role against plaque rupture by stabilizing the fibrous cap. In a unique form of CAD known as plaque erosion VSMCs play a greater role than inflammatory cells in plaque formation (Arbustini, Dal Bello et al., 1999). VSMC dedifferentiation and proliferation contribute not only to atherosclerosis, but also complicate revascularization procedures, promoting restenosis post-angioplasty and intimal hyperplasia at anastomoses in bypass grafts. The molecular mechanisms maintaining or perturbing VSMC differentiation are poorly understood.

The highly conserved Wnt signaling pathway regulates fundamental processes in embryonic development including cell fate and has been implicated in carcinogenesis(Clevers & Nusse, 2012). Recently, Wnt signaling has been implicated in regulation of VSMC differentiation(Srivastava, Zhang et al., 2015). We have previously shown that mutations in the gene encoding Wnt-coreceptor LRP6 underlie autosomal dominant CAD and T2DM in humans (OMIM: ADCADII). Mice with the CAD-linked *LRP6* mutation (*LRP6*^*R611C*^) exhibit coronary and aortic intimal hyperplasia, even in the absence of mechanical vascular injury, largely accounted for by excess VSMC proliferation in the absence of excess inflammation (Srivastava et al., 2015). Disease pathway analysis in *LRP6*^*R611C*^ mice revealed that enhanced noncanonical Wnt signaling enhances PDGF signaling and promotes VSMC dedifferentiation and proliferation and is associated with reduced expression of the Wnt-regulated transcription factor TCF7L2. Wnt3a administration to the *LRP6*^*R611C*^ mice reduced PDGF signaling activities, normalized TCF7L2 expression, and rescued the vascular phenotype. These findings raised the possibility that TCF7L2 may play a causal role in acquisition and maintenance of vascular smooth muscle differentiation. This is an extremely relevant question as common variants (SNP) in the gene encoding *TCF7L2* are considered the strongest genetic risk factors for type 2 diabetes by multiple GWAS, and are associated with risk of CAD (Muendlein, Saely et al., 2011) and its severity (Sousa, Marquezine et al., 2009) in subjects with T2DM. This notion, however, is in sharp contrast with the previous findings that Wnt/TCF7L2 promotes cell cycle activation and proliferation during early embryogenesis and in adult stem cells (Grant, Thorleifsson et al., 2006, He, Kernogitski et al., 2016, Muendlein et al., 2011). Additionally, whether the disease-associated *TCF7L2* common variants cause gain or loss of function has remained controversial (Wang, Xiao et al., 2002). Thus, we embarked on comprehensive studies in mice that either globally overexpress or are haploinsufficient for TCF7L2 to 1) verify the causal role and 2) the mechanisms of action of TCF7L2 in regulation of VSMC plasticity and 3) to establish that TCF7L2-driven VSMC differentiation plays a protective role to counteract the aberrant Wnt signaling and CAD due to the LRP6^R^611^C^ mutation. It should be emphasized that a germline LRP6^R^611^C^ variant promotes human CAD. TCF7L2 GWAS risk alleles are also germline variants and hence their associated traits may arise from their global effects. Therefore, we employ global mouse overexpression or haploinsufficient models to investigate the roles of TCF7L2 as a translationally relevant approach.

## Results

### TCF7L2 expression modulates post-injury intimal hyperplasia

Vascular injury results in VSMC loss of differentiation and their subsequent proliferation (Bennett, Sinha et al., 2016). We first examined the TCF7L2 protein levels in wildtype mice with or without carotid artery wire injury by using immunofluorescent (IF) staining. WT mice carotid arteries were stained with a TCF7L2-specific antibody before and after wire injury. VSMCs in WT carotid artery exhibited robust nuclear staining of TCF7L2 at baseline, which was reduced 3 weeks after wire injury (Fig. 1 A).

**Fig. 1.**
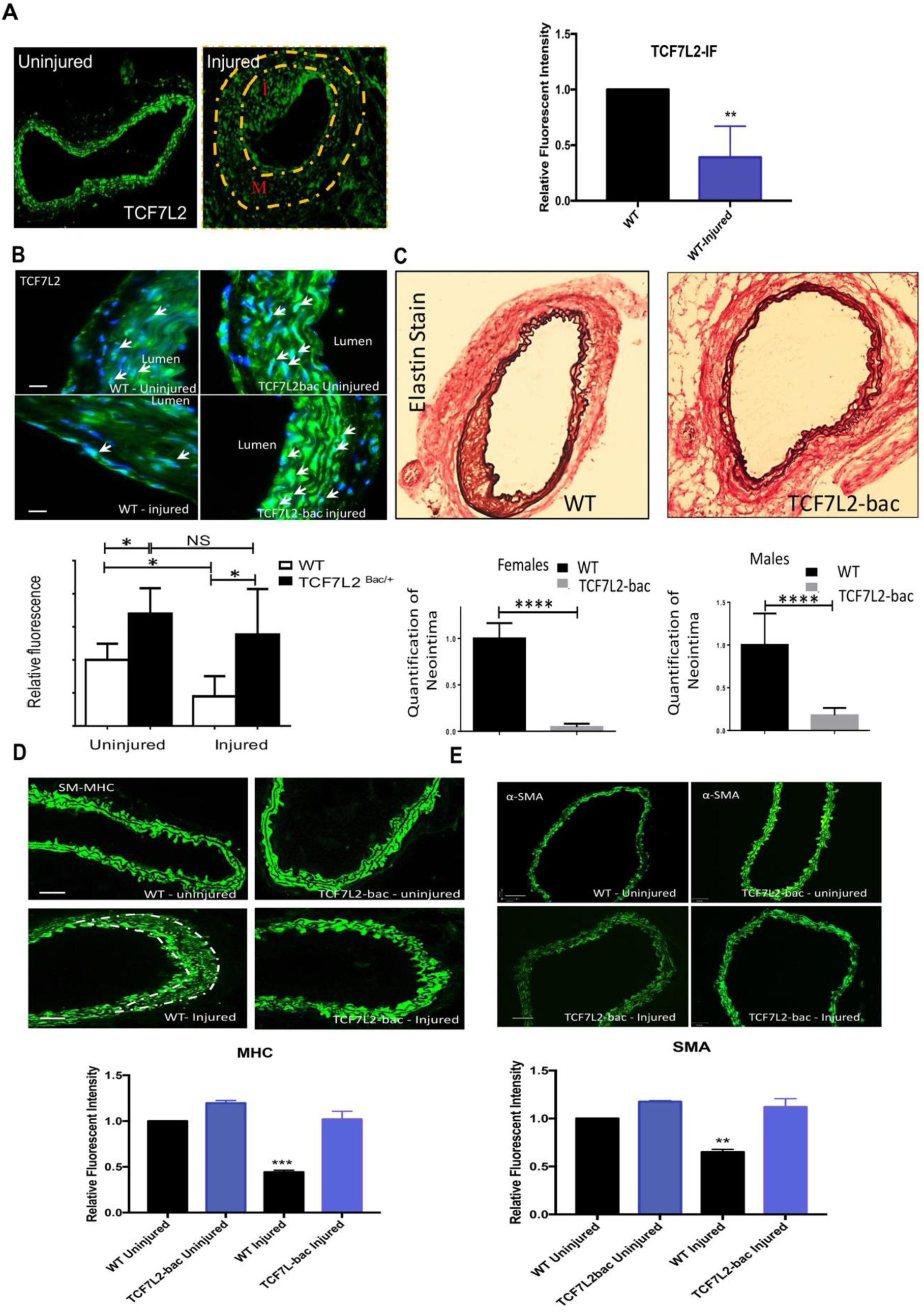
(A-E) Response to mechanical injury in TCF7L2 overexpressing and haploinsufficient mice. (1A) IF staining of TCF7L2 in WT mouse carotid at baseline and post guide wire injury (relative fluorescent intensities shown next to the figure) (1B) TCF7L2 levels in heterozygote overexpressing (TCF7L2-bac) and WT mice before and after injury (relative fluorescent intensities shown underneath). (1C) Neointima formation and EVG staining in carotid arteries of WT and TCF7L2-bac mice post guide wire injury. Quantifications of neointima for both female and male WT, and TCF7L2-bac mice post carotid guide wire injury are shown underneath corresponding figures. (1D-E) immunofluorescent assessment of TCF7L2 SM-MHC and α-SMA (both in green) in carotids of TCF7L2-bac, and WT mice, before and after guide wire injury. The relative fluorescent intensities shown underneath. The yellow dotted lined in 1A demarcate the neointima. *. **, ***, **** denote significance with p value<0.05, <0.01, <0.001and <0.0001, respectively. Quantifications of IF intensity are average of eight sections per mice (7 mice in each group). Scale bars, 16 μm.

To examine the relationship between TCF7L2 levels and VSMC phenotype, we backcrossed BALB/c humanized TCF7L2 overexpressing BAC transgenic mice (transgenic mouse lines harboring engineered enhancer-trapped human bac RP11-466I19 spanning the entire T2D-associated interval recombined with a full-length mouse Tcf7l2 cDNA) and heterozygote knockout mice (generated using zinc finger nuclease technology targeted to the constitutive exon 11 encoding the High Mobility Group box DNA-binding domain of the protein)with C57BL6 mice, until we generated N10 (10 generation) heterozygote overexpressing (TCF7L2-bac) and heterozygous knockout (TCF7L2^+/-^) mice. *Tcf7l2*^*-/-*^ were born in Mendelian ratio but died within ∼24 h of birth. The reason for the backcrossing was 1) LRP6 mice used in the rescue experiments are of C57BL6 background and 2) BALB/c mice are resistant to atherosclerosis (Pei, Wang et al., 2006).

Uninjured TCF7L2-bac mice exhibited > 1.7-fold higher levels of mostly nuclear TCF7L2 in VSMCs compared to WT mice by IF staining (Fig.1B). We then examined the effect of TCF7L2 expression in response to mechanical injury on neointima formation using IF staining and confocal microscopy. As expected, the injury resulted in reduced TCF7L2 in both WT and TCF7L2-bac mice. TCF7L2 levels were, however, 2.5-fold higher in TCF7L2-bac mice vs. wildtype littermates after the wire injury by IF staining (Fig. 1B). Strikingly, TCF7L2-bac male and female mice were both protected against neointima formation post injury compared to same gender wildtype littermates (Fig. 1C). Reduced TCF7L2 IF staining after carotid injury in wildtype mice was associated with marked reduction in the level of contractile proteins SM-MHC (Fig.1D) and α-SMA (Fig.1E), indicating loss of differentiation of VSMC. Conversely, the levels of contractile proteins after the injury were by far higher in TCF7L2-bac vs. WT mice carotids (Fig. 1D, 1E).

We then compared the effect of carotid wire injury on neointima formation in TCF7L2^+/-^ mice compared to their WT littermates. In contrast to TCF7L2-bac mice, TCF7L2^+/-^ mice showed marked reduction of TCF7L2 by IF staining and higher neointima formation post injury compared to wildtype littermates (2A and 2B). Accordingly, the fall in TCF7L2 expression after injury in TCF7L2^+/-^ mice was accompanied by a decline in the levels of contractile proteins SM-MHC and α-SMA in VSMC assessed by IF staining (Fig. 2C, 2D).

**Fig. 2.**
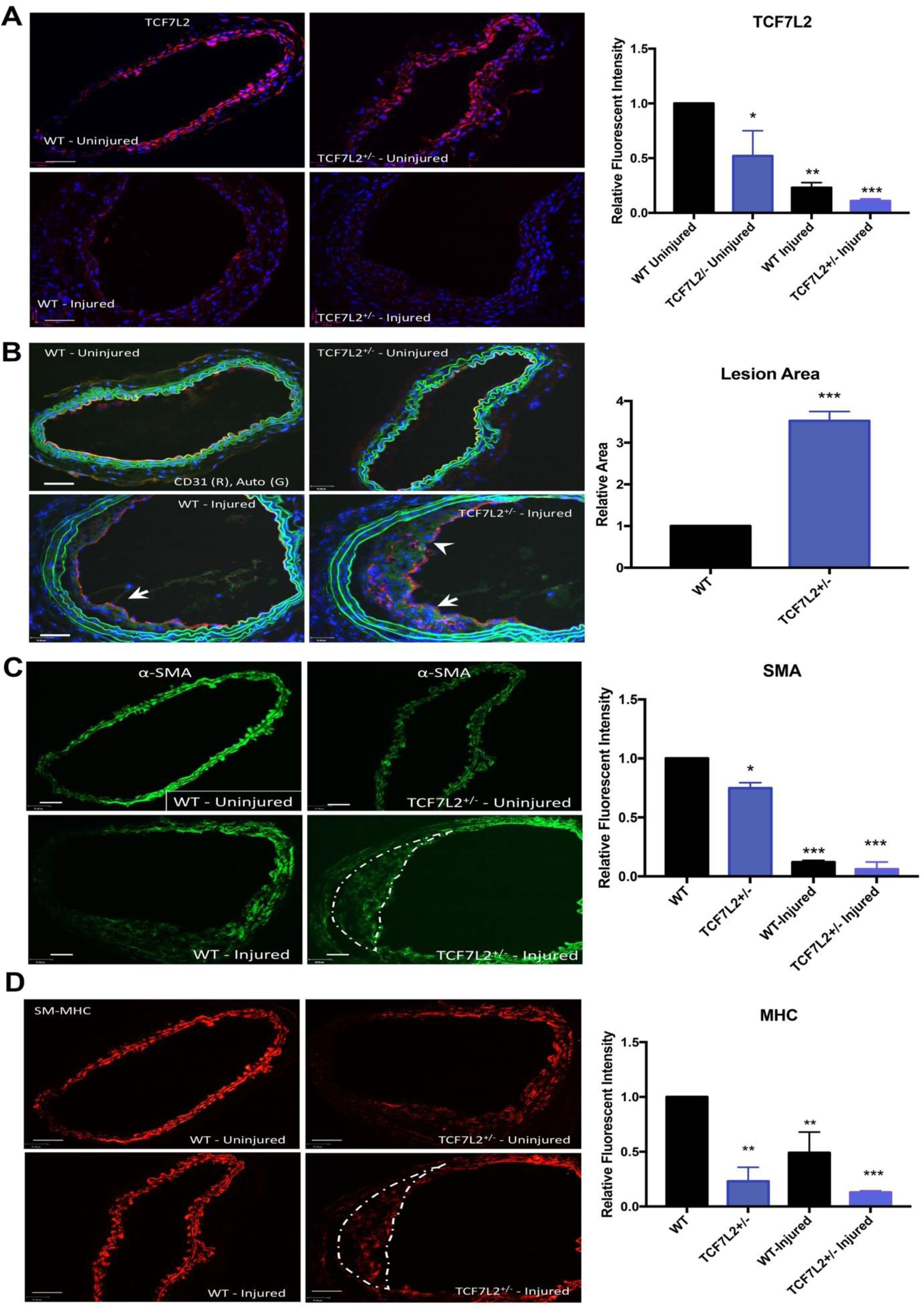
Response to mechanical injury in TCF7L2^+/-^ mice. (2A) IF staining of carotid for TCF7L2 in TCF7L2+/- and WT mice before and after carotid wire injury(2B) Neointima formation and its quantification in carotid arteries of WT and TCF7L2+/- mice post guide wire injury. (2C-D) Immunofluorescent assessment of TCF7L2 SM-MHC and α- SMA in carotids of TCF7L2+/- and WT mice, before and after guide wire injury. Quantifications are shown next to corresponding figures. *. **, *** denote significance with p value<0.05, <0.01, and <0.001, respectively. Quantifications of IF intensity are average of eight sections per mice (7 mice in each group). Scale bars, 16 μm.

### TCF7L2 abrogated VSMC migration and PDGF-stimulated proliferation *in vitro*

Intimal hyperplasia in response to injury is associated with VSMC migration and proliferation. To demonstrate that the protection against intimal hyperplasia in TCF7L2-bac mice is an intrinsic effect of the VSMCs we isolated wildtype and TCF7L2-bac aortic VSMCs and carried out migration and proliferation assays. In *vitro* scratch assay showed that TCF7L2-bac aortic VSMC cultured with FBS migrated considerably slower than WT aortic VSMC (Fig. 3A). PDGF signaling has an established role in triggering VSMC proliferation(Wilson, Mai et al., 1993) (Marmur, Poon et al., 1992). A proliferation assay with and without PDGF–BB stimulation for 24 h was carried out in isolated aortic WT, TCF7L2-bac and TCF7L2^+/-^ VSMC. TCF7L2-bac aortic VSMC showed lower BrdU incorporation and cell count after PDGF–BB stimulation compared with WT aortic VSMC (Fig. 3B, quantifications shown underneath). TCF7L2^+/-^ VSMC had expectedly greater BrdU incorporation and cell number compared to wildtype VSMC after PDGF–BB stimulation (Fig. 3C, quantifications shown underneath). Taken together, these data suggest that TCF7L2 inhibits cell migration and PDGF-dependent cell proliferation.

**Fig. 3.**
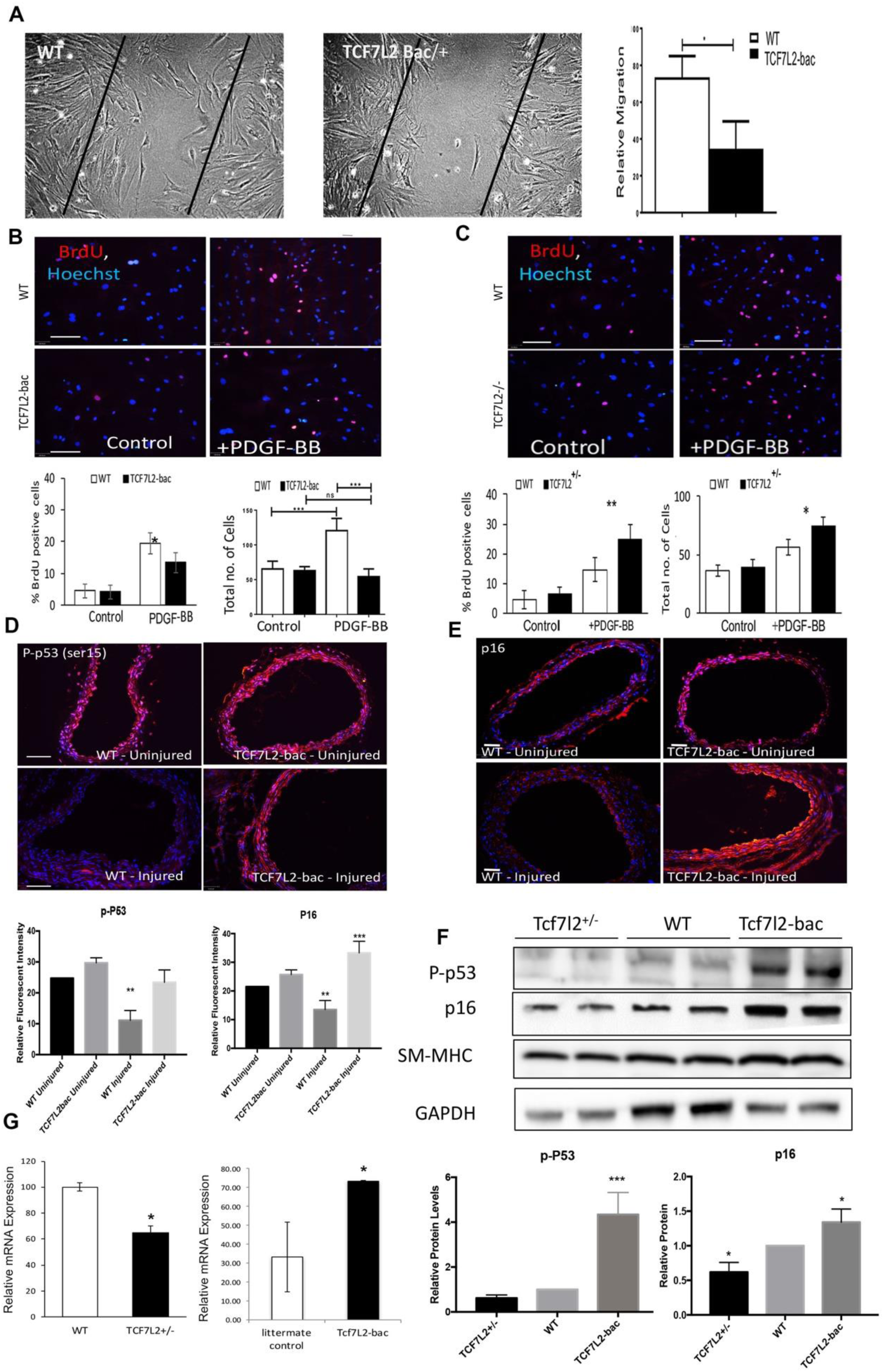
Cell migration and proliferation in TCF7L2-bac and TCF7L2^+/-^ mice VSMCs. (A) *In vitro* scratch assay demonstrating slower TCF7L2-bac VSMC migration as compared to WT VSMC (the quantification shown next to the figure) (B-C) Proliferation assay with BrdU incorporation and cell count analysis with and without PDGF-BB stimulation for 24 hrs in TCF7L2-bac and TCF7L2^+/-^ VSMC, respectively (quantification shown left underneath) (D-E) IF staining of carotid artery for p-P53, and p16 protein in WT and TCF7L2-bac mice before and after injury(quantification shown underneath Fig. D). Proteins levels of p-P53, p16 and SM-MHC in WT, TCF7L2-bac, and TCF7L2^+/-^ mice carotid lysates by Western blot analysis. *, ** and *** denote significance with p values <0.05, <0.01, <0.005 respectively. Quantification are average of 4 experiments per condition. Scale bar, 16 μm.

### TCF7L2 regulates cell Cycle Checkpoints

TP53 (p53) and p16 have been implicated in preventing entry into mitosis as part of cell cycle checkpoint mechanisms. P16 is a downstream target of p53 (Stark & Taylor, 2004, Stark & Taylor, 2006) and is also implicated in G2 arrest during cellular senescence (Gire & Dulic, 2015). We examined p-p53 and p16 levels in WT, TCF7L2^+/-^ and TCF7L2-bac VSMC by immunohistochemistry. Both cell cycle inhibitors were downregulated in wildtype VSMCs but not in TCF7L2-bac VSMC after vascular injury compared to baseline (Fig. 3D-E, quantification underneath 3D). Accordingly, immunofluorescent analysis showed lower p-p53 levels in TCF7L2^+/-^ vs. wildtype VSMCs (Fig. S1A). Western blot analysis confirmed higher baseline p- p53 and p16 protein levels in TCF7L2-bac and lower levels of these proteins in TCF7L2^+/-^ carotids compared to their wildtype littermates (Fig. 3F, quantifications underneath the figure).

It has been previously shown that TCF7L2 binds the p53 promoter and transcriptionally regulates this gene (Zhou, Zhang et al., 2012). We therefore examined the p53 mRNA levels, which were dramatically lower in TCF7L2^+/-^ and higher in TCF7L2-bac compared to wildtype VSMCs (Fig.3G). Notably, this effect of TCF7L2 in VSMCs is in contrast with its effect in proliferating pancreatic beta cells in which TCF7L2 represses p53 expression (Zhou et al., 2012).

### Association between TCF7L2 and GATA6 in VSMC

GATA6 has been recognized as a key regulator of VSMC differentiation (Wada, Hasegawa et al., 2000). This transcription factor promotes VSMC differentiation by stimulating the expression of contractile proteins and inhibiting cell cycle progression (Perlman, Suzuki et al., 1998). The shared function of TCF7L2 and GATA6 with respect to promoting the expression of contractile proteins prompted an examination into possible interaction between the two proteins. We compared the GATA6 protein levels of WT and TCF7L2-bac mice carotid arteries before and after wire injury by IF staining. Baseline GATA6 levels were modestly higher in TCF7L2-bac compared to WT mice VSMCs (Fig.4A). GATA6 levels were almost undetectable after injury in wildtype mice, but GATA6 levels in TCF7L2-bac mice post-injury were considerably higher compared to wildtype mice. We then co-stained for TCF7L2 and GATA6 and examined their subcellular localization in VSMCs of the tunica media of WT mice with and without carotid artery injury using confocal microscopy. The results showed that in quiescent arteries, TCF7L2 and GATA6 colocalize within the nucleus in VSMC, but their expression levels drastically dropped after the injury and with the nuclear exclusion of TCF7L2 they no longer colocalized (Fig.4B).

These findings raised the question whether TCF7L2 regulates GATA6 expression. GATA6 transcripts are down-regulated in response to mitogens (Suzuki, Evans et al., 1996). PDGF is a major mitogen that has an established role in inducing loss of differentiation and proliferation of VSMCs post-injury (Holycross, Blank et al., 1992, Reusch, Wagdy et al., 1996) and is inhibited by Wnt activation (Keramati, Singh et al., 2011, Srivastava et al., 2015). To examine if TCF7L2 transcriptionally regulates GATA6, we treated wildtype and TCF7L2-bac VSMC with or without PDGF-BB for 24 hours and measured the GATA6 and SM-MHC mRNA expression levels by qRT-PCR. Baseline GATA6 mRNA levels were higher in TCF7L2-bac vs. wildtype VSMC. The mRNA expression levels of TCF7L2 and GATA-6 were both dramatically reduced after PDGF-BB stimulation in WT VSMC. As anticipated, there was a concurrent decline in SM-MHC (aka. *MYH11*) mRNA expression (Figure 4C). Notably, PDGF-induced repression of TCF7L2, GATA6, and SM-MHC mRNA levels was significantly attenuated in TCF7L2-bac aortic VSMC (Fig. 4C).

**Fig. 4.**
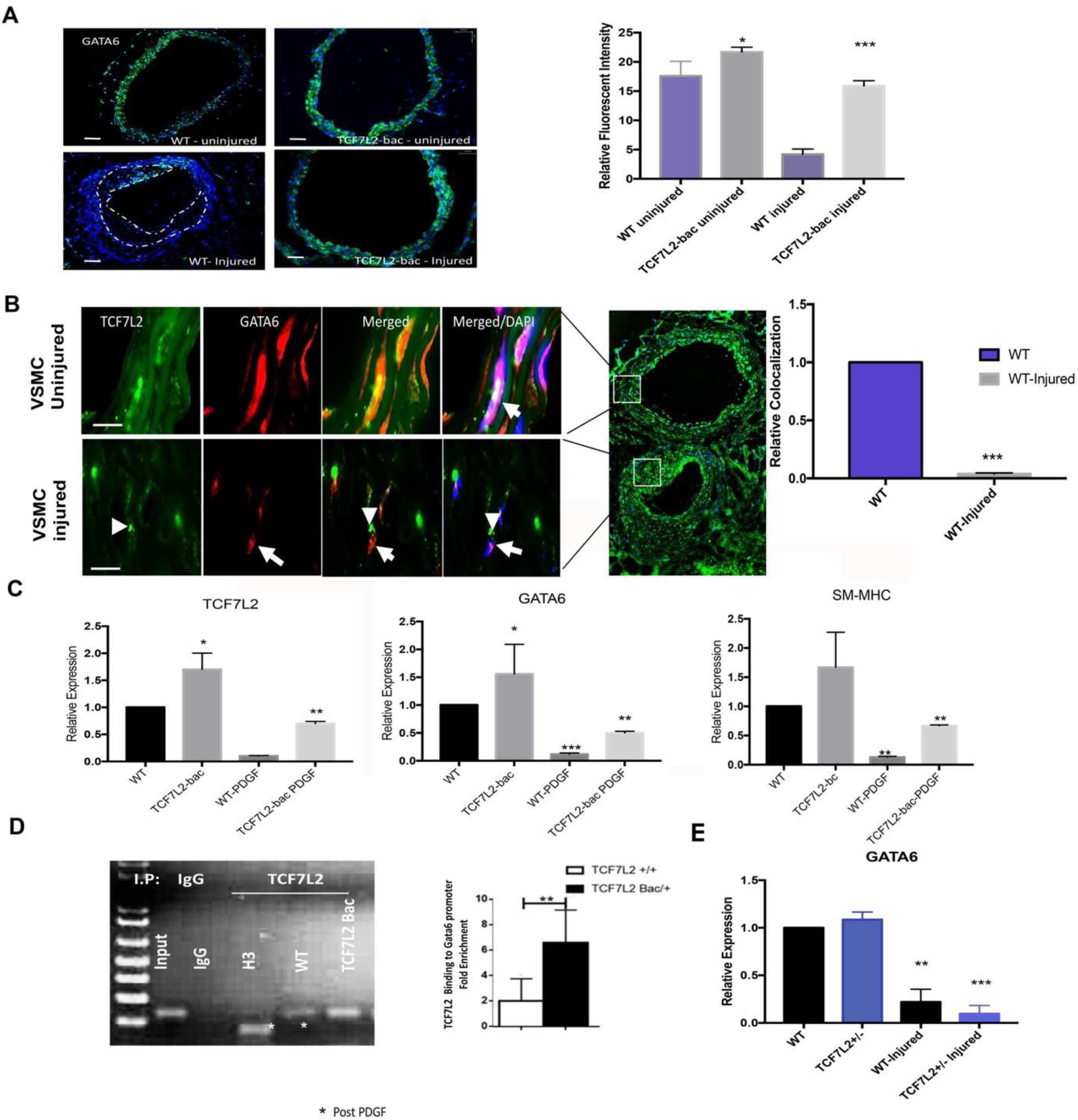
GATA6 Expression, Localization and transcriptional regulation. (3A) IF staining of GATA6 in WT and TCF7L2-bac mice carotids, before and after guide wire injury (quantification shown next to the figure). (3B) Subcellular localization of TCF7L2 and GATA6 in tunica media of injured and uninjured carotid artery of WT mice. Colocalization of TCF7L2 and GATA6 in the nucleus and lack of thereof after injury, arrows and arrow heads show GATA6 and TCF7L2 respectively (3C) TCF7L2, GATA6 and SM-MHC mRNA expression in TCF7L2-bac vs. WT VSMC, with and without PDGFBB stimulation (D) ChIP assay demonstrating TCF7L2 binding to GATA6 promoter; its quantification shown next to the figure. (E) GATA6 mRNA expression in WT and TCF7L2^+/-^ mice aortic lysate, before and after guide wire injury. *. **, *** denote significance with p value<0.05, <0.01, and <0.001, respectively. Quantifications of IF intensity are average of eight sections per mice (7 mice in each group). Scale bars, 16 μm.

The GATA6 promoter has multiple (T-C-A-A-A-G or C-T-T-T-G-A) motifs as putative sites for TCF7L2 binding. We, therefore, carried out a ChIP assay to examine TCF7L2 binding to the GATA6 promoter by amplifying the region of the mouse GATA6 promoter bearing the putative TCF7L2 binding sites using specific primers. The assay revealed specific binding of TCF7L2 to the GATA6 promoter (Figure 4D). GATA6 promoter binding by TCF7L2 was expectedly stronger in TCF7L2-bac vs. wildtype VSMC (Figure 4D). We then compared GATA6 mRNA levels between wildtype and TCF7L2+/- carotid artery lysate before and after injury. The GATA6 mRNA levels were not different between the wildtype and TCF7L2+/- mice carotids at baseline. TCF7L2-bac the GATA6 mRNA and protein levels assayed by IF were dramatically reduced after injury in both wildtype and TCF7L2+/- mice carotids (Fig. 4 E and Fig. S1B). These findings suggested that TCF7L2 protects VSMCs against PDGF-mediated transcriptional inhibition of GATA6, but is not required to maintain it under normal condition. Importantly, the effects of TCF7L2 on GATA6 mRNA levels were modest and did not entirely account for the more robust effects observed at the protein level in vivo. This prompted an investigation into posttranscriptional regulation of GATA6 by TCF7L2.

### TCF7L2 is protective against PDGF-dependent turnover of GATA6

Activation of the PDGF downstream effector JNK has been shown to promote the proteolytic degradation of GATA6 (Ushijima & Maeda, 2012). JNK activator anisomycin has been shown to efficiently export nuclear GATA-6 into the cytoplasm and initiates its degradation by proteasomes, and this effect is inhibited by JNK inhibitor SP600125. We had previously shown that LRP6/Wnt inhibits PDGF signaling by promoting PDGFβR ubiquitination and diminishing JNK signaling. Subsequent *in vitro* an *in vivo* investigations had found that impaired Wnt signaling results in increased activation of PDGF–signaling (Srivastava et al., 2015), which is normalized by systemic Wnt3a administration *in vivo* and by TCF7L2 overexpression in VSMCs *in vitro* (Srivastava et al., 2015). Thus, we hypothesized that TCF7L2 may stabilize GATA6 protein by inhibiting PDGF/JNK activation. To examine this hypothesis, we measured GATA6 protein levels in VSMC of WT and TCF7L2-bac mice with and without PDGF-BB treatment after inhibiting protein synthesis by cycloheximide. The analysis showed greater GATA6 protein levels in TCF7L2-bac vs. WT VSMCs. PDGF–BB dramatically reduced GATA6 protein levels in wildtype VSMC within 15 minutes of stimulation (Fig.5A), whereas GATA6 was only marginally reduced in TCF7L2-bac and after 30 minutes of stimulation. This robust and reproducible finding implicated TCF7L2 in stabilization of GATA6 against PDGF–BB.

We then examined the effect of TCF7L2 overexpression on PDGF signaling in WT vs. TCF7L2-bac VSMCs stimulated with PDGF–BB. There was significantly lower total and phosphorylated PDGFRβ in TCF7L2-bac vs. wildtype VSMC after PDGF–BB stimulation (Fig.5B). Consequently, there was reduced activation of its PDGF downstream effectors, JNK/SAPK/ERK in TCF7L2-bac compared to wildtype VSMC (Fig. 5B). These findings identify TCF7L2 inhibition of PDGF/JNK as one of the mechanisms for GATA6 stabilization.

**Fig. 5.**
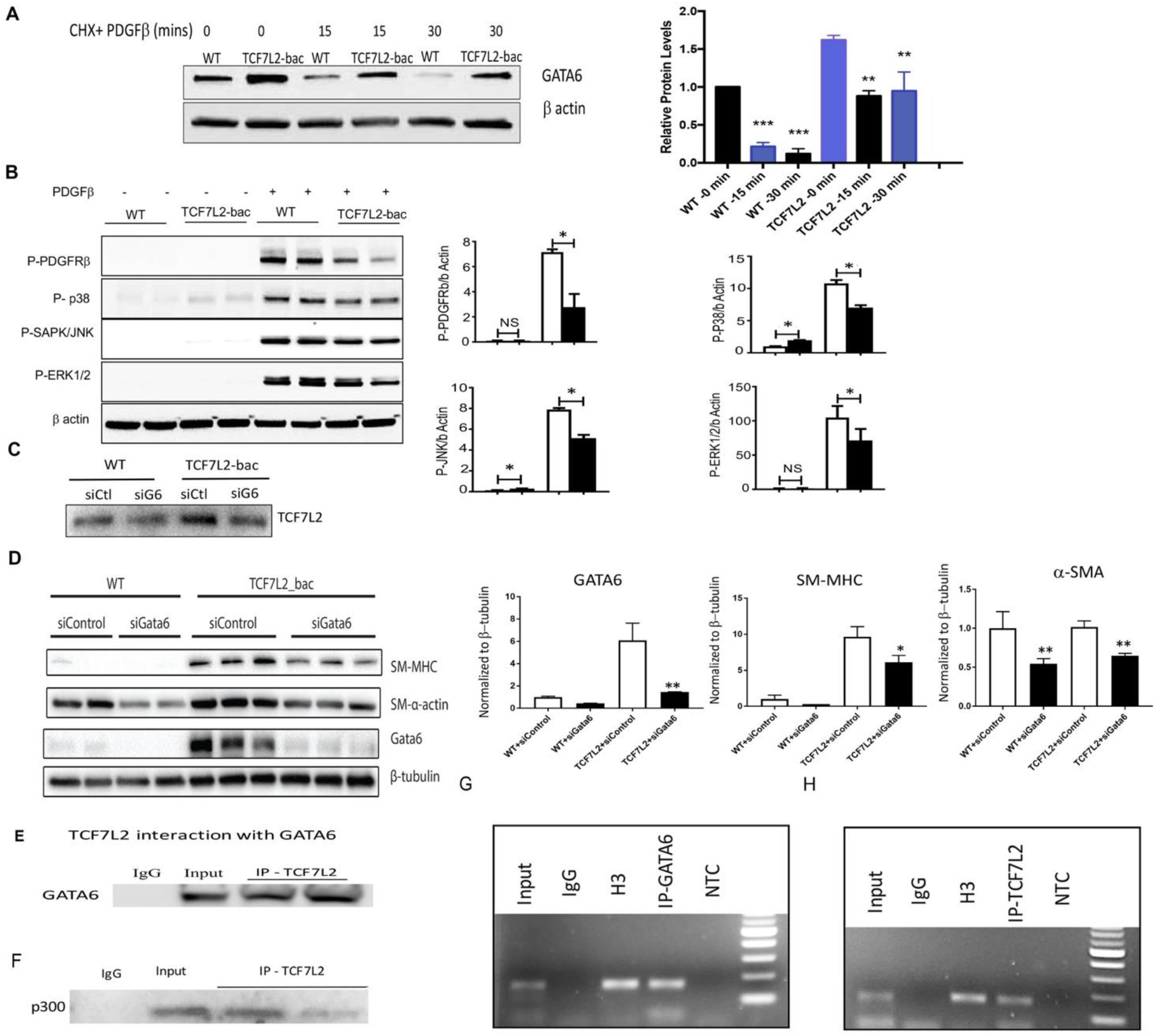
TCF7L2 stabilizes GATA6 against PDGF, forms a complex with it and jointly regulates SM-MHC transcription. (4A) Time course of GATA6 protein levels following PDGF-BB stimulation in CHX pretreated WT and TCF7L2-bac VSMC by western blot, quantifications shown underneath the figure, quantifications shown next to the figure (4B) PDGFRβ expression and phosphorylation and activation of its downstream effectors, RhoA, p38, ERK1/2 and JNK/SAPK in WT and TCF7L2-bac VSMC upon PDGF-BB stimulation, quantifications shown next to the figure (4C) GATA6 knockdown by siRNA and (4D) its effect on the protein levels of MYH11 and SMA in WT and TCF7L2-bac VSMC, quantifications shown next to the figure (3E) Immunoprecipitation of TCF7L2 and GATA6 in WT and TCF7L2-bac VSMC lysates, TCF7L2 specific antibody was used for pulldown, followed by Western blot analysis for GATA6 (3F) Immunocoprecipitation of TCF7L2 and p300 in WT and TCF7L2-bac VSMC lysates, TCF7L2 specific antibody was used for pulldown, followed by Western blot analysis for p300 (4G, H) ChIP assay demonstrating bindings of TCF7L2 and GATA6 to SM-MHC promoter using ChIP-grade TCF7L2-specific and GATA6-specific antibody, respectively. *, ** and *** denote significance with p values <0.05, <0.01, <0.001 respectively. Quantification are average of 4 experiments per mice (7 mice in each group).

**Fig. 6.**
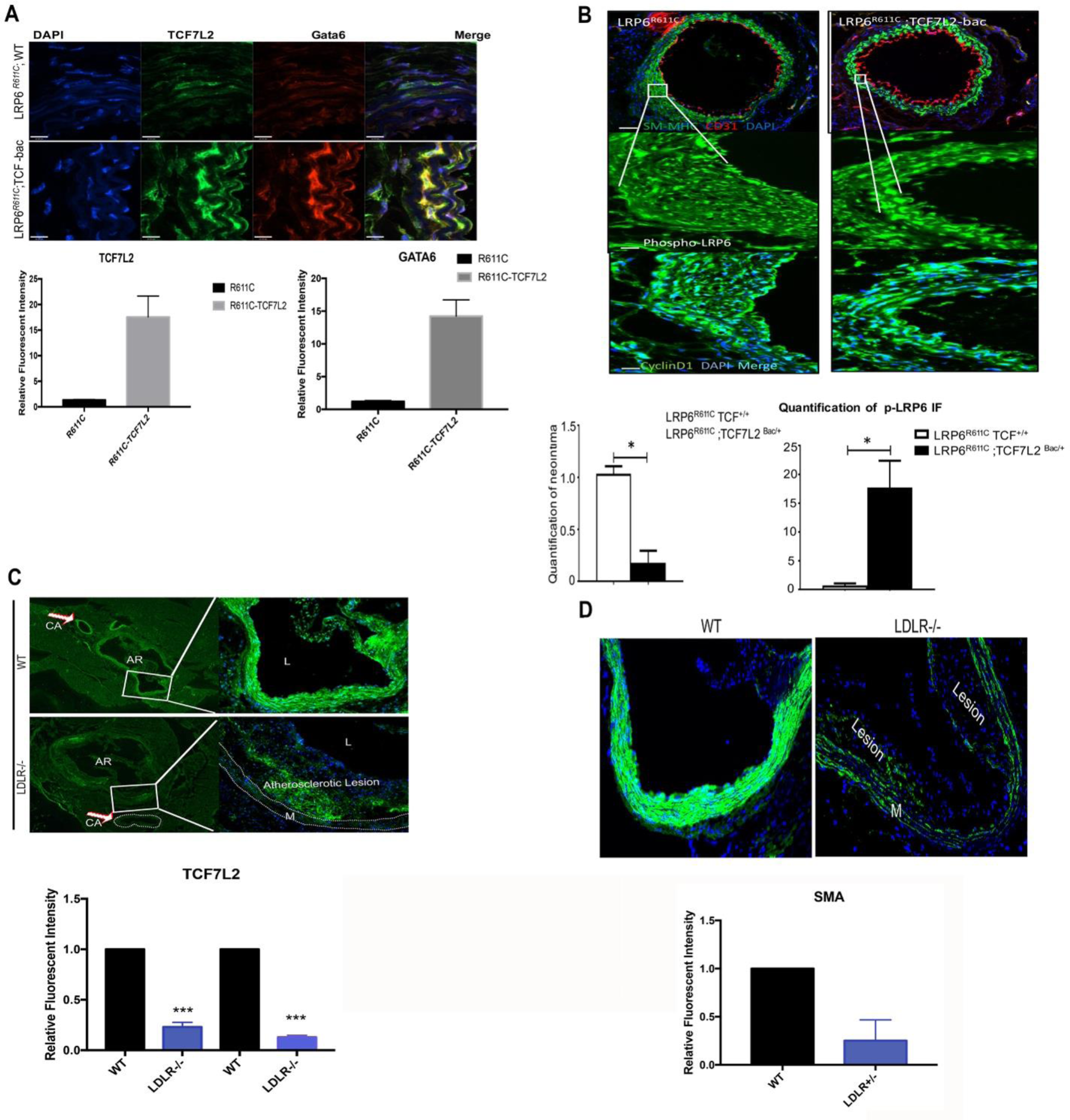
TCF7L2 Overexpression rescues post carotid injury neointima formation in *LRP6*^*R611C*^ mice. (A) relative intensity of TCF7L2 and GATA6 proteins in *LRP6*^*R611C*^ and *LRP6*^*R611C*^; TCF7L2-bac mice carotids by IF (B) Neointima formation and IF staining of carotid artery post injury for SM-MHC, p-LRP6 and cyclinD1 expression in *LRP6*^*R611C*^ vs. *LRP6*^*R611C*^; TCF7L2-bac mice(C-D) TCF7L2 and α-SMA levels with quantifications (underneath corresponding figures) in the aortic root and coronary arteries of the LDLR^−/-^ vs. WT mice on high fat diet.AR denotes aortic root. CA denotes coronary artery. A section of aortic root is enlarged. *, ** and *** denote significance with p values <0.05, <0.01, <0.001 respectively. Scale bars, 16 μm.

### TCF7L2 and GATA6 complex formation and binding of SM-MHC promoter

To verify if GATA6 mediates the TCF7L2 regulation of SM-MHC we knocked down GATA6 in WT and TCF7L2-bac VSMCs by siRNA. GATA6 protein levels were markedly increased in the TCF7L2-bac and were knocked down by about 80% by GATA6 siRNA (Fig.5C). Consequently, SM-MHC protein levels were significantly reduced in TCF7L2-bac VSMCs compared to control siRNA (siControl, quantifications are shown next to the figure) (Fig.5D). SM-MHC protein levels continued to be higher in TCF7L2-bac compared to WT cells, either due to incomplete knockdown of GATA6 or GATA6-independent regulation of SM-MHC by TCF7L2. Taken together, these findings suggested that TCF7L2 regulates SM-MHC via GATA6-dependent and potentially GATA6-independent mechanisms.

Given that TCF7L2 and GATA6 colocalize in the nucleus, have similar functions and exhibit similar responses to perturbations, the possibility that they may have a combinatorial effect on SM-MHC transcription was raised. Thus, we examined if TCF7L2 and GATA6 form a complex. A pull-down assay using TCF7L2-specific antibody followed by Western blot analysis using GATA6-specific antibody showed that TCF7L2 and GATA6 coimmunoprecipitate (Fig. 5E). GATA6 has been shown to form a complex with p300 to regulate transcription of SM-MHC in a combinatorial fashion (Wada et al., 2000). An immunoprecipitation assay in VSMC showed that TCF7L2 also forms a complex with p300 (Fig. 5F), suggesting a potential combinatorial interaction between GATA6, TCF7L2 and p300. We then carried out an *in-silico* screening of genes encoding VSMC contractile proteins for putative consensus TCF7L2 binding motifs. Interestingly, several consensus binding sequences for TCF7L2 and GATA6 were found either in tandem or overlapping by one or 2 nucleotides. We focused on one such consensus (CTTTGATAA) that lies between untranslated exon 1 and exon2 in the open chromatin region of *SM-MHC* gene as defined by DNase I hypersensitivity (Fig. S1C, shown in the ascending aorta). We carried out two independent ChIP assays using ChIP-grade antibodies specific for TCF7L2 or GATA6. The assays revealed that both proteins bind to a region containing this motif in mouse VSMC (Fig. 5G-H). Overall, these findings indicate that TCF7L2 and GATA6 are both implicated in transcriptional regulation of SM-MHC gene and suggest a potential combinatorial interaction between the two in this process.

### TCF7L2 Overexpression rescues post carotid injury neointima formation in *LRP6*^*R611C*^mice

We had previously shown that impaired LRP6 activity in *LRP6* mutant mice (*LRP6*^*R611C*^) leads to enhanced non-canonical Wnt signaling, and a subsequent increase in TCF7L2 turn over and reduced SM-MHC protein levels (Srivastava et al., 2015). These cascades of events culminate in excessive intimal hyperplasia. To 1) establish a causal link between reduced TCF7L2 expression, VSMC dedifferentiation and intimal hyperplasia, and 2) demonstrate that TCF7L2-mediated VSMC differentiation and reduced proliferation is protective against intimal hyperplasia, a rescue experiment in *LRP6*^*R611C*^ mice was carried out. We crossbred TCF7L2-bac onto *LRP6*^*R611C*^ mice and examined the effect of TCF7L2 overexpression on GATA6 levels and SM-MHC in carotid injury neointima formation. Compared to *LRP6*^*R611C*^ mice, *LRP6*^*R611C*^; TCF7L2-bac mice exhibited higher TCF7L2 and GATA6 protein levels (Fig.6A, quantification underneath). We then examined whether TCF7L2 overexpression rescues carotid artery intimal hyperplasia following guidewire injury in *LRP6*^*R611C*^ mice. As expected, TCF7L2 overexpression had protective effects against neointima formation in *LRP6*^*R611C*^; TCF7L2-bac (Fig.6B, quantifications underneath). Furthermore, increased GATA6 levels in *LRP6*^*R611C*^; TCF7L2-bac mice were associated with increased staining for SM-MHC compared to *LRP6*^*R611C*^ mice. Of note, the overexpression of TCF7L2 was also associated with increased levels of activated (phosphorylated) LRP6 phosphorylation (Fig.6B), which may suggest a positive feedback mechanism. The exact molecular mechanisms of this phosphorylation has not been elaborated (Davidson, Shen et al., 2009).

### TCF7L2 expression in atherosclerosis model

We had previously shown that TCF7L2 is highly expressed in VSMCs (Srivastava et al., 2015). However, its significance in atherosclerosis process had not been studied. We assessed TCF7L2 protein levels in the aortic root and coronary arteries of low-density receptor knockout (LDLR^−/-^) vs. WT mice. Strikingly, TCF7L2 levels were also lower in the aortic root and coronary arteries of LDLR^−/-^ vs. WT mice on high fat diet (Fig.6C). These changes were also associated with reduced α-SMA levels in VSMC of LDLR^−/-^ mice compared to WT mice (Fig.6D).

### TCF7L2 risk alleles are associated with reduced gene expression in the aorta

Common variants of TCF7L2 gene have been the strongest signal in GWAS of T2DM and age-related endophenotypes (Grant et al., 2006) (p< 8 x10^−17^) and are associated with risk of CAD (Muendlein et al., 2011) and its severity (Sousa et al., 2009) in subjects with T2DM. Due to small effect size of these variants, however, the disease mechanisms have remained unclear. Delineating TCF7L2 function in regulation of VSMCs is critical for understanding the pathogenesis of arterial disease in type 2 diabetes. Analyses of GTEX project data revealed that DM2 and CAD-associated TCF7L2 minor alleles (i.e. rs4506565, Fig. S1D) are in disequilibrium with the expression QTLs variants of TCF7L2 (i.e. rs6585200 and rs6585201, p<1.5e-10, NES - 0.31; Fig. S1E). Based on GTEX data QTLs variants of TCF7L2 are most significantly associated with reduced TCF7L2 expression in the aorta and tibial artery (Fig. S1F, S1G, http://www.gtexportal.org/home/eqtls/byGene?geneId=tcf7l2&tissueName=All). Taken together, these findings in human atherosclerosis and LDLR knockout mice indicate the relevance of our findings in the TCF7L2 mouse models and the significance of TCF7L2 expression levels for normal arterial function.

## Discussion

Mutations in Wnt-coreceptor LRP6 regulated proteins have been associated with coronary artery disease (CAD) (Mani, Radhakrishnan et al., 2007). The causality of LRP6 mutation has been established in a mouse model of human mutation that exhibits severe obstructive CAD, characterized by excessive proliferation of poorly differentiated VSMCs (Srivastava et al., 2015). The roles of downstream effectors of Wnt/LRP6 are poorly understood and, in some cases, controversial. Here, we examine in detail the role of TCF7L2 in VSMC phenotype and report that it is an important regulator of VSMC differentiation. This function of TCF7L2 as we demonstrate is achieved, in part, through stabilization of GATA6. Mitogens secreted after injury are known to reduce GATA6 transcripts (Suzuki et al., 1996). In addition, GATA6 is proteosomally degraded by JNK, which is an effector of PDGF, one of the most the most potent mitogens in VSMCs (Ushijima & Maeda, 2012). We demonstrate that TCF7L2 reduces total PDGFRβ protein and diminishes JNK activation in VSMC. We then show that both TCF7L2 and GATA6 are required for MYH11 transcription, form a complex together, and both bind an open chromatin region in the MYH11 promoter containing a tandem TCF7L2/GATA motif. Our findings elucidate a previously unrecognized molecular interaction between these transcription factors and identify TCF7L2 as a novel regulator of GATA6-dependent VSMC phenotypic switching.

During early embryogenesis and in adult stem cells TCF7L2 activates cell cycle and promotes cell proliferation (Wang et al., 2002). This has led to the conventional wisdom that TCF7L2 promotes VSMC proliferation. Our investigation showed that TCF7L2 inhibits VSMCs proliferation by upregulating p-P53 and its downstream effector P16. Strikingly, GATA6 has been also shown to increase the expression of p53 and induce growth inhibition in VSMCs (Perlman et al., 1998).

Our study raises several important questions that require future investigations. These include how does injury or PDFG-BB downregulate TCF7L2, and are the TCF7L2 effects cell-autonomous or non-cell-autonomous? Such questions may be better addressed in different models such as tissue-specific knockout mice. One should bear in mind that tissue–specific knockouts may result in completely different traits than those observed in humans simply because the TCF7L2 GWAS-variants are germline polymorphisms and the associated traits may result from complex interaction between different organs and tissues that express these genetic variants. In this regard, the generation of TCF7L2 global haploinsufficient or overexpressing mice was an essential step to account for the potential large-scale effects of TCF7L2 variants.

We identify TCF7L2 as a key effector of Wnt signaling based on its ability to rescue the intimal hyperplasia phenotype of the LRP6^R^611^C^ mutant in injury models. As these mutant mice are also prone to coronary and aortic intimal hyperplasia characterized by medial and intimal thickening in the absence of significant macrophage accumulation, it would be of interest to determine the role of TCF7L2 in atherogenesis versus fibrous cap formation in this model in future studies.

In summary, our study unravels the role of TCF7L2 in regulation of VSMC plasticity and protection against arterial disease. Our earlier work had uncovered the role of TCF7L2 in regulation of plasma lipids (Go, Srivastava et al., 2014) and glucose (Singh, De Aguiar et al., 2013). Collectively, our findings underscore the important role of TCF7L2 as a link between metabolic traits and arterial disease and identifies it as an important target for development of drugs to combat T2DM, metabolic syndrome and its vascular complications.

## Experimental Procedures

### Animals

Animal procedures were as per approved protocol of Yale University Institutional Animal Care and Use Committee. TCF7L2 overexpression (TCF7L2-bac under mouse TCF7L2 promoter) and TCF7L2 heterozygous knockout (TCF7L2 +/-) mice were obtained in collaboration from Dr. Marcelo Nobrega’s laboratory at The University of Chicago, Chicago, IL (Savic, Ye et al., 2011). These mice were then backcrossed to C57Bl6 mice (N>10). Wildtype littermates were used as controls. Generation of homozygous *LRP6*^*R611C*^ and *LRP6*^*R611C*^; LDLR^−/-^ mice were previously described (Go et al., 2014). Generation of *LRP6*^*R611C/R611C*^; TCF7L2-bac was done by crossing *LRP6*^*R611C/R611C*^ to C57Bl6 backcrossed TCF7L2-bac mice. *LRP6*^*R611C/R611C*^ homozygous mice are referred to as either as *LRP6*^*R611C*^ in the text or *R611C* or *RC* mice in the figures. LDLR knockout mice were obtained from Jackson laboratory, Bar Harbor, ME. All mice were fed ad libitum and housed at constant ambient temperature in 12 h light, 12 h dark cycle. Mice were examined at age 6-8 weeks. For high-cholesterol diet studies 6 weeks old mice were fed high-cholesterol diet (40% fat, 1.25% cholesterol, and 0.5% cholic acid) ad libitum (Research Diets) for 4 or 6 weeks.

### Chemicals and Antibodies

Protease inhibitor cocktail (P8340), phosphatase inhibitor cocktail (P2850), cycloheximide, MG132 and BrdU were purchased from Sigma-Aldrich. Cell lysis buffer (9803) and antibodies for β-actin, PDGFRβ, pPDGFRβ (y751/y771), p-LRP6 vimentin and KLF4, p-RhoA, p-JNK, p-ERK1/2, cyclin D1, ChiP assay kit were all purchased from Cell Signaling Technology. DMEM, fetal bovine serum (FBS), penicillin streptomycin cocktail, Trypsin-EDTA solution, and TRIzol were purchased from GIBCO/Invitrogen; polyvinylidene fluoride membranes from Bio-Rad Laboratories; Antibodies for TCF7L2, and protein A/G agarose gel were purchased from Santa Cruz Biotechnology. Antibodies for α-SMA, and SM-MHC were purchased from Abcam, CD31 antibody from BD PharMingen, and secondary fluorescence tagged antibodies and propidium iodide from Invitrogen.

### Immunofluorescence

Immunofluorescence staining was performed on 5-μm frozen sections, and fluorescence was measured using Nikon Eclipse80i or Zeiss 4 laser Confocal microscope using same laser output, gain, and offset for each set of antibodies tested. Images were quantified with Image J.

### Western blotting

Whole-cell lysates of primary VSMCs were separated by electrophoresis, transferred to PVDF membrane, and probed using target primary antibodies followed by appropriate HRP-conjugated secondary antibodies. Blots were visualized using chemiluminescence reagents, imaged with Bio-Rad gel doc system, and quantified with Image J software. Real-Time PCR.

### VSMC Isolation and Culture

Isolation of aortic VSMCs was carried out as previously described (Ray, Leach et al., 2001). VSMCs were maintained in DMEM (4.5g/l glucose, glutamine, and 100 mg/l sodium pyruvate) supplemented with 20% FCS, 100 U ml-1 penicillin, and 100 μg ml-1 streptomycin. For phosphorylation studies, cells were starved for 3 h in 0.2% FBS containing DMEM and were treated as follows: PDGF-BB at 10 ng/ml for 15 min. For mRNA studies, the cells were starved overnight in DMEM containing 0.2% FBS and treated with PDGF-BB (10 ng/ml) for 24 hr. For protein half-life measurements, VSMC were pretreated with cycloheximide (CHX) (100ug/mL) for 30 minutes and then incubated with or without PDGFBB (10ng/mL). Cells were harvested at 0, 15 and 30 minute intervals, cell lysates were subjected to western blotting and the GATA6 protein was quantified using Image J. For proteosomal degradation studies, VSMC were pretreated with MG132 (1μM) for 1h followed by PDGFBB (10 ng/mL) stimulation. The cell lysates were immunoprecipitated using GATA6 antibody, followed by western blotting and probing with ubiquitin antibody.

### ChIP Assay

Chromatin immunoprecipitation (ChIP) assay was performed according to the manufacturer’s instructions (Cell Signaling SimpleChIP^®^ Enzymatic Chromatin IP Kit (Agarose Beads) #9002). Briefly, the chromatin/DNA protein complexes were prepared from mouse aortic smooth muscle cells. Chemical crosslinking of DNA proteins was carried out using 1% formaldehyde for 10 min at room temperature and followed by addition of glycine solution. Cells were scraped into cold PBS containing Halt cocktail proteinase inhibitor. The cell suspension was centrifuged and the pellet was lysed and nuclei was digested using micrococcal nuclease to digest DNA to a length of approximately 200–1,000 bp. Supernatant containing the digested chromatin was incubated with appropriate ChIP-grade TCF7L2 antibody (sc-8631; Santa Cruz Biotechnology) for immunoprecipitation overnight at 4°C with rotation, followed by ChIP-grade protein A/G agarose beads and incubation for 1h at 4°C with rotation. Anti-H3 antibody and RPL30 primers provided in the kit were used as a positive control for assay technique and reagent integrity. The agarose resin was washed using buffers supplied with the kit. The eluted DNA was purified and analyzed by PCR to determine the binding of TCF7L2 (TCF7L2 binding consensus: (A/T) (A/T) CAAAG) to Gata6 promoter. The following primers were used to amplify the binding region: forward 5’-GTCTTCGACGCCTAGCTTCA-3’ and reverse 5’-CTAAATTTGGCGTCCTGGCTG-3’. Real-time PCR amplification was performed using iQ SYBR Green Supermix (Bio-Rad) and Eppendorf Mastercycler RealPlex2. Similar protocol used to amplify TCF/GATA6 binding site (ctttgataa) in MHY11 intron2 with the primers 5’-TGCCTGGGTTTCATTCTGTG −3’ and 5’-GGCCCTCACCTCTCCATATC −3’.

### Carotid Artery Guide Wire Injury

The carotid artery guide wire injury was performed as previously described (Wang, Zhang et al., 2009) Three weeks post-injury, mice were euthanized and injured carotid arteries were excised from the arteriotomy site of external left carotid artery, including the internal left carotid artery and approximately 1 cm of left common carotid artery. Similarly, right common carotids were harvested and used as uninjured controls. The arteries were embedded in OCT; serial tissue sections (5 μm) were obtained from left and right common carotid arteries, starting at the bifurcation (to external and internal carotids); and immunofluorescence (IF), IHC, and morphometric analyses were performed. Neointima formation was measured in ten sections (50 μm apart) using images obtained by a bright-field microscope and quantified using ImageJ software (NIH).

### *In vitro* scratch wound healing assay

Primary VSMC from WT and TCF7L2-bac aorta were grown to ∼70-80% confluency in 12 well plates in complete growth medium. The monolayer was scratched across the center of the well gently and slowly using tip of sterile 1mL pipette tips and holding the tip perpendicular to the bottom of the well. After scratching, the wells were gently washed twice with culture medium to remove detached cells and filled with fresh culture medium. Microscope images were taken to note the initial scratch size and the areas were marked. The cells were then left undisturbed for 18h and images were taken again. The gap distance covered was quantitatively evaluated using Image J software.

### *In vitro* BrdU staining

VSMC were grown to 70-80% confluency and starved overnight (DMEM with 0.2% FBS). Next cells were stimulated with or without PDGFBB (10ng/mL) followed by propidium iodide for nuclear staining for 24 hours. Then cells were pulsed with BrdU (10uM/mL) for 4 hours at 37°C. BrdU solution was removed and washed with PBS (3 times, 2 minutes each). The cells were fixed with 1mL of 3.7% formaldehyde for 15 minutes at room temperature (RT) and washed with PBS (3 times, 2 minutes each). Next, the cells were permeabilized with 1ml of 0.1% Triton^®^ X-100 in PBS for 20 minutes at RT, followed by 1mL of 1N HCl for 20 minutes at RT and finally neutralized with 0.1 M sodium borate buffer pH 8.5 for 30 minutes at RT, washed with 0.1% Triton^®^ X-100 in PBS (3 times, 2 minutes each). The cells were incubated overnight with anti-BrdU antibody at 4°C, washed with 0.1% Triton^®^ X-100 in PBS (3 times, 2 minutes each) followed by incubation with fluorescently labeled secondary antibody for 1h at RT. The cells were washed with PBS and imaged using fluorescence microspore.

### Real Time PCR

Total RNA was isolated from primary VSMC culture using TRIzol, and cDNA was generated using the High Capacity cDNA Reverse Transcription Kit (Applied Biosystems) according to the manufacturer’s instructions. Real-time PCR amplification was performed using specific primers and iQ SYBR Green Supermix in Eppendorf Mastercycler RealPlex2. Reactions were performed in quadruple with a β-actin internal control. Relative quantification of mRNA levels was expressed as fold increase relative to the control. The following mouse primer sequences were used for qRT-PCR:

- Gata6 Forward: ATGCGGTCTCTACAGCAAGATGA
- Gata6 Reverse: CGCCATAAGGTAGTGGTTGTGG
- TCF7l2 Forward: CGCTGACAGTCAACGCATCTATG
- TCF7L2 Reverse: GGAGGATTCCTGCTTGACTGTC
- SM-MHC Forward: GCAACTACAGGCTGAGAGGAAG
- SM-MHC Reverse: TCAGCCGTGACCTTCTCTAGCT
- βactin Forward: CATTGCTGACAGGATGCAGAAGG
- βactin Reverse: TGCTGGAAGGTGGACAGTGAGG

## Statistical Analyses

Guide wire injury studies were done using at least 10 mice per group. All *in vitro* studies were carried out in three independent experiments in triplicate. Fluorescence and area measurements were done using Image J software (NIH). Preparation of graphs and all statistical analyses of results by 2-tailed Student’s *t* test were carried out using GraphPad Prism 6 Project software (GraphPad). p < 0.05 was considered significant. Data are presented as mean ± SD.

## Acknowledgement

The Genotype-Tissue Expression (GTEx) Project was supported by the Common Fund of the Office of the Director of the National Institutes of Health, and by NCI, NHGRI, NHLBI, NIDA, NIMH, and NINDS. The data used for the analyses described in this manuscript were obtained from: [http://www.gtexportal.org/home/?geneId=TCF7L2] the GTEx Portal on 10/28/2015. TCF7L2-bac transgenic and heterozygote TCF7L2 knockout mice were generous gifts from Dr. Marcelo Nobrega from University of Chicago. The study was funded by R35 HL135767 grant form NIH to A.M.

---

**Suppl. fig. 1.**
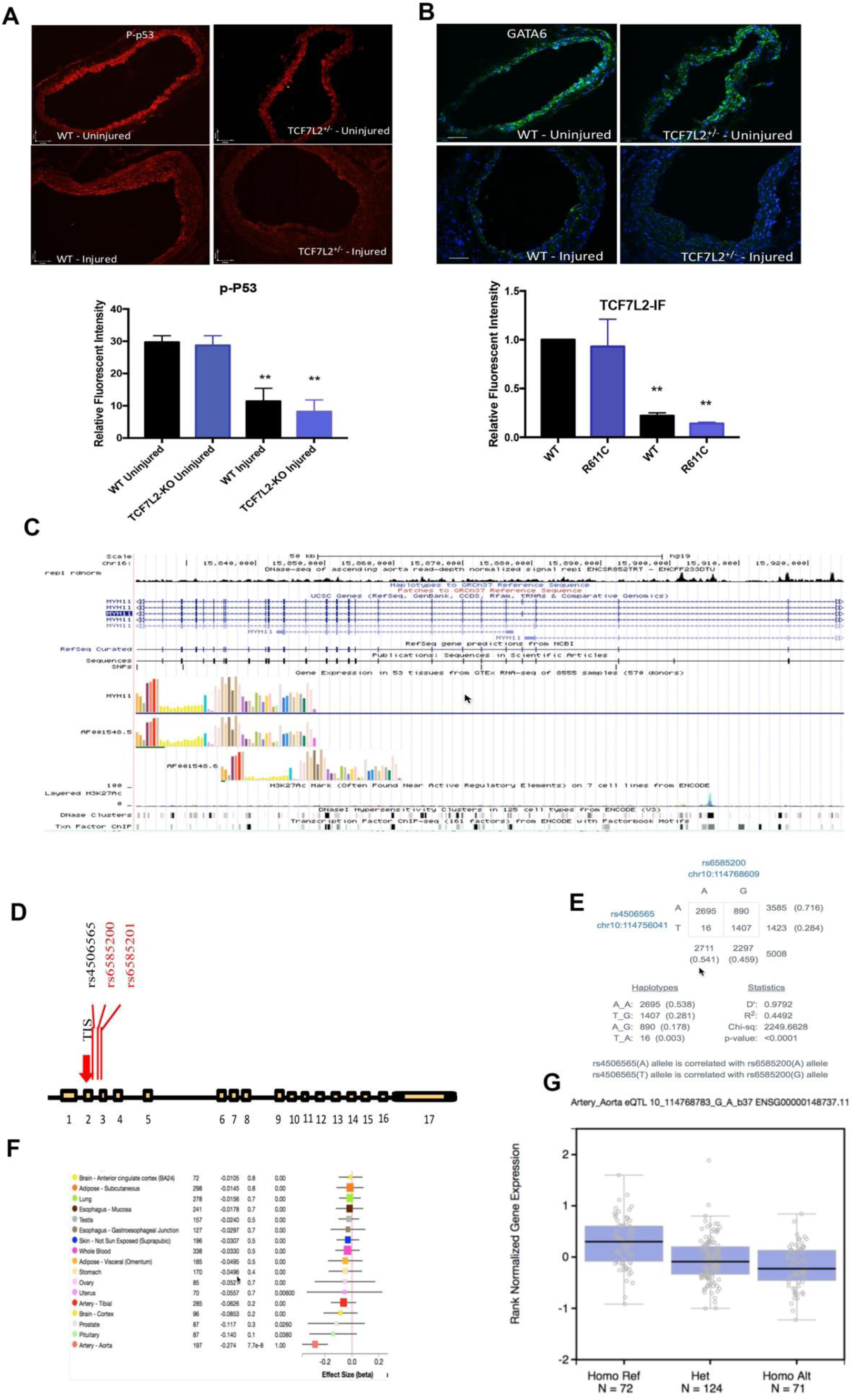
A) Immunofluorescent staining for p53 in TCF7L2^+/-^ vs. wildtype VSMCs, B) IF staining of carotid for GATA6 in WT and TCF7L2^+/-^ mice, before and after guide wire injury and their quantification (C) DNase I hypersensitivity sites in MYH11 locus in the heart. The first peaks in the intron corresponds to GATA6/TCF7L2 binding site (from The ENCODE (Encyclopedia of DNA Elements) https://www.encodeproject.org (D)The position of GWAS SNPS: rs7901695and rs7903146 and the eQTL SNP rs7895340 within TCF7L2 gene (E) disequilibrium between rs7903146 and the eQTL SNP rs7895340 (C, D) single tissue eQTL p-values for the rs7895340 and the effect size adopted from GTEX database (http://www.gtexportal.org/home/eqtls/byGene?geneId=tcf7l2&tissueName=All). Scale bars, 16 μm.

